# Plasma ST6Gal1 is Dispensable for IgG Sialylation

**DOI:** 10.1101/2022.02.01.478679

**Authors:** Douglas M Oswald, Sylvain D Lehoux, Julie Y Zhou, Leandre M Glendenning, Richard D Cummings, Brian A Cobb

**Affiliations:** Case Western Reserve University School of Medicine, Department of Pathology, Cleveland, OH, USA; Beth Israel Deaconess Medical Center, Harvard Medical School Center for Glycoscience, National Center for Functional Glycomics, Boston, MA, USA

**Keywords:** ST6Gal1, glycobiology, IgG, sialylation, hepatocyte, plasma, sialic acid

## Abstract

The glycosylation of IgG has attracted increased attention due to the impact of N-glycan modifications at N297 on IgG function, acting primarily through modulation of Fc domain conformation and Fcγ receptor binding affinities and signaling. However, the mechanisms regulating IgG glycosylation and especially α2,6-sialylation of its N-glycan remain poorly understood. We observed previously that IgG is normally sialylated in mice with B cells lacking the sialyltransferase ST6Gal1. This supported the hypothesis that IgG may be sialylated outside of B cells, perhaps through the action of hepatocyte-released plasma ST6Gal1. Here we demonstrate that this model is incorrect. Animals lacking hepatocyte expressed ST6Gal1 retain normal IgG α2,6-sialylation, despite the lack of detectable ST6Gal1 in plasma. Moreover, we confirmed that B cells were not a redundant source of IgG sialylation. Thus, while α2,6-sialylation is lacking in IgG from mice with germline ablation of ST6Gal1, IgG α2,6-sialylation is normal in mice lacking ST6Gal1 in either hepatocytes or B cells. These results indicate that IgG α2,6-sialylation arises after release from a B cell, but is not dependent on plasma-localized ST6Gal1 activity.

## Introduction

Research beginning in the early 1980s has firmly established the links between changing biological and disease states with alterations in IgG glycosylation (reviewed in (1)). Human and mouse IgG contains a single and conserved N-linked glycan at N297 of the Fc domain in each heavy chain, which is primarily a biantennary complex N-glycan varying in of GlcNAc bisection, galactosylation, core fucosylation, and terminal α2,6-linked sialylation (2). A number of early studies characterized IgG glycosylation in the context of rheumatoid arthritis (RA) and reported a decrease of galactosylation and sialylation associated with disease. Similar studies involving a variety of inflammatory conditions have extended these findings to establish that inflammatory diseases lead to a decrease of galactosylation and sialylation, directly demonstrating that these modifications to the IgG glycan are under regulatory control.

The importance of sialylation of IgG was documented first in 2006, when it was reported that the active portion of high dose intravenous immunoglobulin therapy (IVIg) responsible for its anti-inflammatory properties in autoimmune patients was the α2,6-sialylated glycoforms of IgG (3). Subsequent studies have largely documented the linkage of IgG glycosylation changes to various physiological and pathological states including pregnancy (4,5), tuberculosis (6), HIV (7,8), diabetes (9), kidney dysfunction (10), lupus (11), and others, as has been recently reviewed (1). Mechanistic work has focused on how sialylated IgG reduces inflammation (12,13), but many outstanding questions remain (14,15).

Our recent work has established that the α2,6-sialylation of IgG is independent of the required enzyme ST6Gal1 in B cells (16), using a novel B cell-specific conditional knockout of ST6Gal1 (BcKO). Despite lacking ST6Gal1, these BcKO animals express α2,6-sialylated IgG at levels indistinguishable from wild type (WT) mice. Although there are published data on the relationship of B cell ST6Gal1 expression and IgG sialylation (17), as well as other models of B cell glycan modification (18), it is clear that B cell ST6Gal1 is not required for IgG sialylation (16). Instead, the observations with the BcKO mouse point to a previously suggested possibility (19), based on a growing body of work, that the complex N-glycans of IgG and other circulatory glycoproteins can be modified after they are secreted into the fluid phases of the body – a process called extracellular sialylation.

ST6Gal1 is highly and inducibly expressed in hepatocytes as a membrane-bound enzyme (20); however, due to action by the protease BACE-1, a soluble form of ST6Gal1 can be released into plasma (21,22), where ST6Gal1 retains enzymatic activity (16,23). Coupling our BcKO observations (16) with previous findings linking liver ST6Gal1 with IgG sialylation (16,19), we and others hypothesized that the liver-released ST6Gal1 may be required for the synthesis of sialylated IgG. Indeed, the development of mice lacking the liver-specific P1 region of the ST6Gal1 promoter (ΔP1) was reported to have both reduced circulatory ST6Gal1 activity (23) and reduced IgG sialylation (19). Consistent with this model, we showed that ST6Gal1 expression in the liver is dramatically increased in halos surrounding central veins where hepatocyte proteins and glycoproteins are released into circulation. Finally, there is evidence that human platelet granules may supply the nucleotide-activated sugars that serve as donors for glycosyltransferases within the plasma (24), and our prior findings confirm that murine platelets can also supply CMP-sialic acid for ST6Gal1 activity (16).

However, to conclusively establish whether extracellular sialylation from plasma-localized ST6Gal1 may be contributing to α2,6-sialylation of IgG, we generated a hepatocyte-specific conditional knockout of ST6Gal1 (HcKO) using the albumin-Cre mouse, which we have recently reported (25,26). We observed, as predicted, a complete ablation of α2,6-linked sialic acids on liver-produced circulatory glycoproteins and hepatocyte cell surfaces, but not in other cells or tissues. Here, we describe the impact of hepatocyte ablation of ST6Gal1 on IgG sialylation. Our results indicate that despite the loss of detectable plasma ST6Gal1 in the HcKO mice, IgG sialylation was unchanged compared to WT controls. Additionally, BACE-1 knockout mice also had normal IgG sialylation. Importantly, mice with germ-line loss of ST6Gal1 lacked sialylated IgG, thereby confirming the absolute requirement for ST6Gal1 in IgG sialylation. We further developed a mouse lacking ST6Gal1 in both the hepatocyte and B cell compartments (BHcKO) to determine whether each cell might be able to compensate for the other. IgG sialylation remained unchanged. These findings demonstrate that neither ST6Gal1 in plasma nor in B cells generates IgG sialylation, and suggest that an unidentified compartment outside of these must be necessary to regulate the sialylation of IgG.

## Results

Total plasma glycoprotein glycosylation in WT and HcKO mice was characterized with and without ovalbumin in alum immunization using a plate-based lectin ELISA assay as we have previously described (27). We found that regardless of immunization status, SNA signal was very low in HcKO samples (Fig 1A). Interestingly, MAL-II, WGA, and PHA-E all showed marked increases in naïve HcKO mice compared to WT, but those differences and overall levels were dramatically reduced in both strains upon immunization, despite that the immunized plasma samples were taken 6 weeks after the final boost to allow the mice to return to baseline. Conversely, immunization increased LCA and AAL, both fucose-detecting lectins (Fig 1A). Principal component analysis (PCA) of these data illustrated the major differences between naïve WT and HcKO plasma glycosylation (Fig 1B). Immunization significantly reduced the difference between strains, but the immunized cohort was significantly different from their naïve counterparts (Fig 1B), with the exception that SNA-detectable α2,6-sialylation remained distinct between strains (Fig 1A).

**Figure 1.**
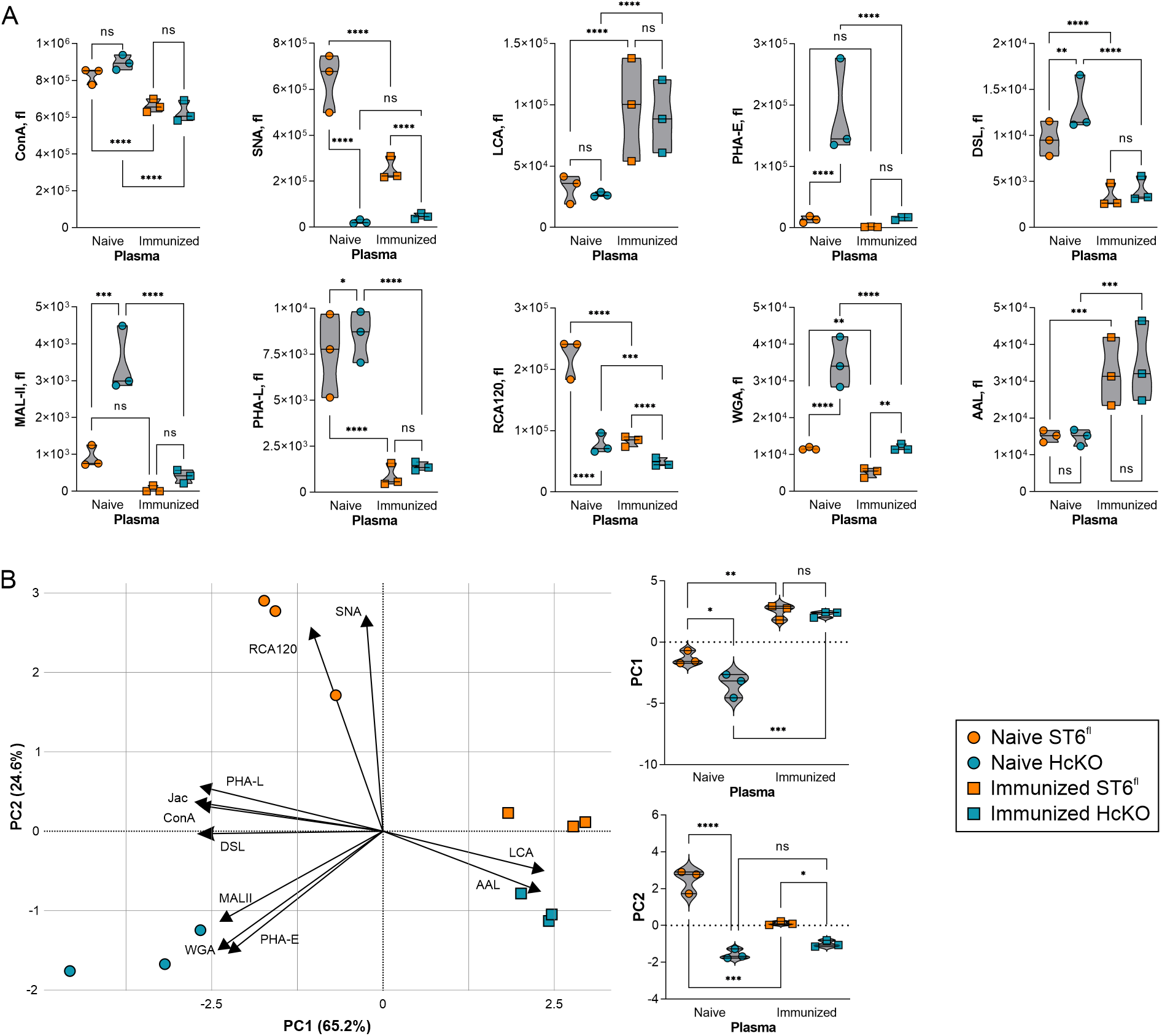
Plasma glycoprotein glycan changes are associated with both hepatocyte ST6Gal1 and immunization. (A) Lectin ELISA of total plasma glycoproteins in resting and immunized WT and HcKO mice 2 weeks after the final booster exposure using an array of lectins revealed a loss of SNA in HcKO samples, and reductions in ConA, SNA, PHA-E, DSL, PHA-L, RCA120, and WGA in WT and HcKO associated with immunization. Increased LCA and AAL was also seen in both strains following immunization. (B) PCA results demonstrated that immunized mouse plasma glycans are more similar to each other than either naïve mouse. PC1 and PC2 scores for each of the four groups show the dominance of immunization in determining overall plasma glycan characteristics. N=3 for each condition and strain.

Next, the IgG response to immunization was quantified. We found that both WT and HcKO mice strongly responded to immunization, although immunized WT mice did responded stronger (Fig 2A). SNA and anti-Fc antibody ELISA on purified IgG from each immunized strain revealed no detectable difference in total sialylation between strains (Fig 2B). The dramatically increased presence of IgG might explain the decreases in SNA and ConA, and increased LCA signal in the total plasma samples of immunized mice (Fig 1A) due to the low sialylation and high fucosylation characteristics of typical IgG N-glycans (2).

**Figure 2.**
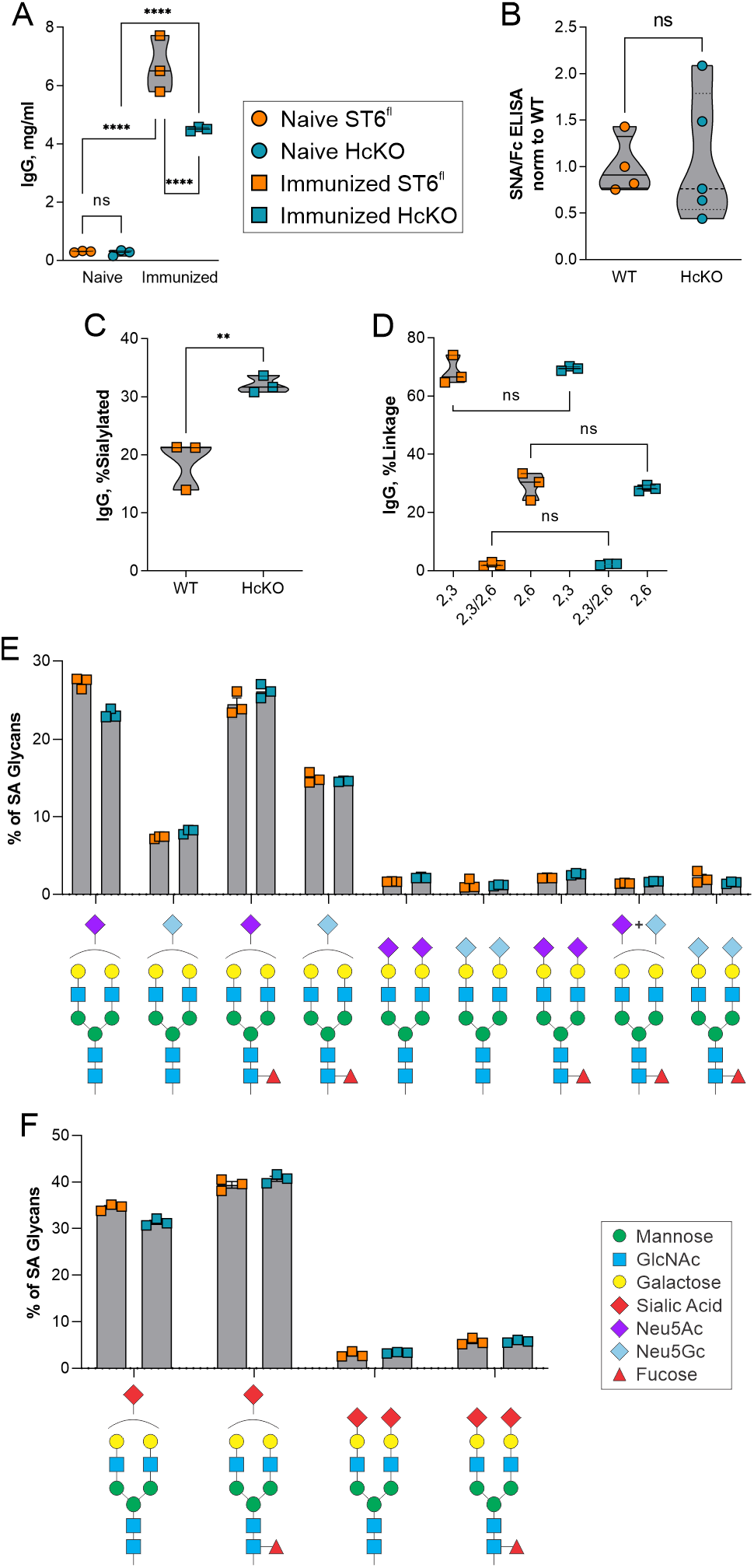
IgG glycan sialylation is consistent between WT and HcKO mice. (A) IgG titers in immunized and non-immunized mice, 2 weeks following the final immunization, showed strong IgG responses in both strains, despite a slightly reduced overall response in HcKO mice. N=3 (B) Lectin ELISA data on purified IgG revealed a lack of difference in α2,6-sialylation between WT and HcKO IgG. N=4-5 (C) Based on mass spectrometry, total IgG sialylated glycan species were increased in HcKO mice. N=3 (D) No difference in linkage distribution of sialic acids was detected. N=3 (E and F) A breakdown of individual sialylated glycan species found by mass spectrometry showed that the distribution of mono-sialylated, di-sialylated, Neu5Ac, Neu5Gc, and fucosylation was indistinguishable between WT and HcKO mice. N=3

To more robustly quantify α2,6-sialylation, we performed mass spectrometry analysis on IgG glycans using 4-(4,6-dimethoxy-1,3,5-triazin-2yl)-4-methylmorpholinium chloride (DMT-MM) activation of the sialic acid carboxylate group, which upon permethylation leads to selective amidation of α2,6-linkages and spontaneous lactonization of α2,3-linkages that are easily differentiated by mass spectrometry due to an added +13 Da on α2,3-linked sialic acids (28). The degree of sialylation was found to be approximately 20 % in WT mice, but over 30 % in HcKO mice (Fig 2C). We found that the majority of sialic acids in murine IgG are present in an α2,3 linkage, whereas α2,6 linkages were a minority, although there was no significant difference between strains (Fig 2D), consistent with prior studies (2). A detailed breakdown of sialylated N-glycans further reveals that the dominant structures were G2S1 with either Neu5Ac or Neu5Gc, and plus or minus core fucose (Fig 2E and 2F). Very few di-sialylated N-glycans were present in either strain on IgG at this time point (i.e. 2 weeks post final boost immunization).

BACE-1 has been shown to be necessary for ST6Gal1 release from the liver (22). Thus, the BACE-1 knockout mouse would be expected to phenocopy HcKO mice in terms of IgG sialylation due to a lack of circulatory ST6Gal1. We obtained plasma from naïve WT and BACE-1 knockout mice from the Jackson Laboratories. IgG concentration was higher in the unimmunized BACE-1 knockout mice (Fig 3A), and the degree of IgG α2,6-sialylation, as measured by SNA binding, was indistinguishable from WT mice from the same facility (Fig 3B). Finally, we confirmed that ST6Gal1 was indeed required for IgG α2,6-sialylation using the germline ST6Gal1 knockout mouse. IgG from these mice lacked SNA-detectable α2,6-sialylation (Fig 3C).

**Figure 3.**
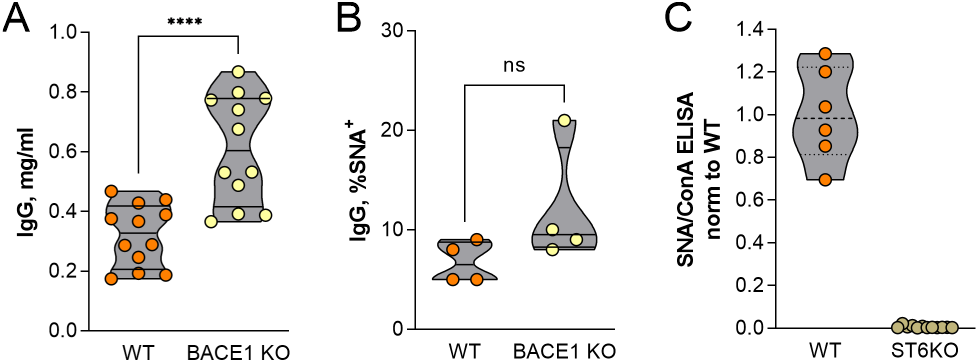
ST6Gal1 but not BACE-1 is necessary for IgG α2,6-sialylation. (A) BACE-1 knockout mice carried modestly but significantly higher IgG titers compared to similarly housed WT control mice. N=12 (B) The degree of α2,6-sialylation of IgG from resting and non-immunized WT and BACE-1 knockout mice, as measured by SNA, was indistinguishable. N=4 (C) Mice with germline knockout of ST6Gal1 lacked detectable IgG α2,6-sialylation. N=6-9

These data demonstrate that hepatocyte-released ST6Gal1 is dispensable for IgG sialylation, suggesting two possibilities. One is that either B cells and hepatocytes (i.e. circulatory ST6Gal1) are redundant pathways and thus can compensate for each other, yielding a lack of IgG sialylation differences in each of the HcKO and BcKO mouse models. The other is that neither of these compartments are necessary for IgG sialylation. In order to differentiate between these two possibilities, we created a mouse in which ST6Gal1 was ablated in both hepatocytes and B cells (BHcKO; Fig 4A). We first confirmed the lack of hepatocyte α2,6-sialylation using confocal microscopy and found a similar pattern of SNA staining in the BHcKO liver (Fig 4B) as we previously reported for the HcKO mouse (25,26). Likewise, B cells were analyzed by flow cytometry with SNA and ECL, and we found the same decrease in SNA and increase in ECL (Fig 4C) as we previously reported for the BcKO mouse (16). These data confirm that the ablation of ST6Gal1 was found in both target tissues as expected.

**Figure 4.**
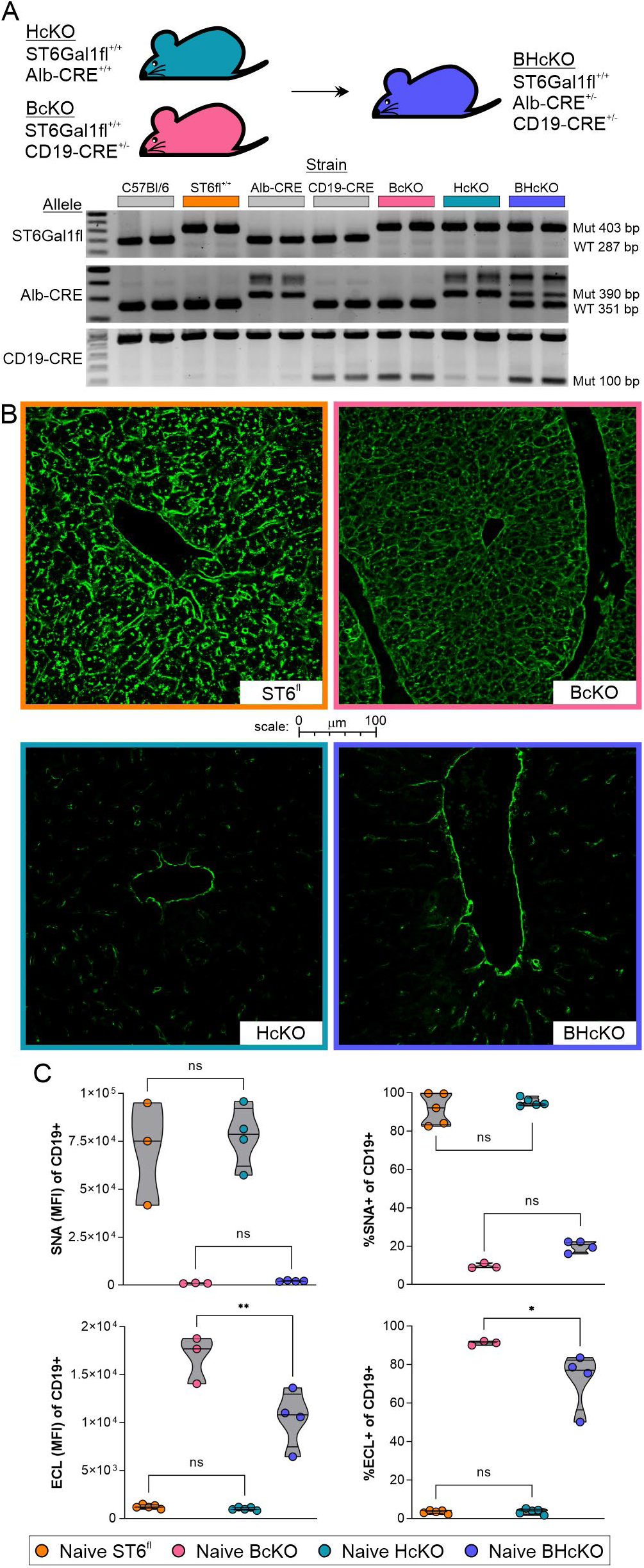
Creation and general characterization of the BHcKO mouse. (A) Scheme, genetics, and validating genotype analysis of the dual-deficient mutant mice. (B) Confocal staining for SNA in the liver revealed the expected deficiency of sialylation in HcKO and BHcKO, but not WT and BcKO tissue; image border color description is indicated in panel A. (C) SNA and ECL lectin flow cytometry analysis of blood CD19^+^ B cells demonstrated the ablation of surface α2,6 sialic acids and increase in terminal galactose respectively on BcKO and BHcKO mice cells, but not in WT and HcKO. N=3-5

Before analyzing the IgG itself, we also confirmed that the HcKO and BHcKO mice lacked ST6Gal1 activity in the plasma using a plate assay as we have previously described (27). We found that both the WT and BcKO strains showed strong ST6Gal1 activity in plasma samples when supplied with exogenous CMP-sialic acid, but neither the HcKO nor BHcKO mice showed detectable activity (Fig 5A). Likewise, we confirmed that plasma-localized ST6Gal1 retains activity *in vivo* by injecting CMP-sialic acid via the tail vein and monitoring for IgG sialylation. Although the total IgG concentration did not change, the degree of IgG sialylation increased (Fig 5B), suggesting that the enzyme in circulation can function if given adequate donor, but that it is dispensable in maintaining IgG sialylation with or without immunization.

**Figure 5.**
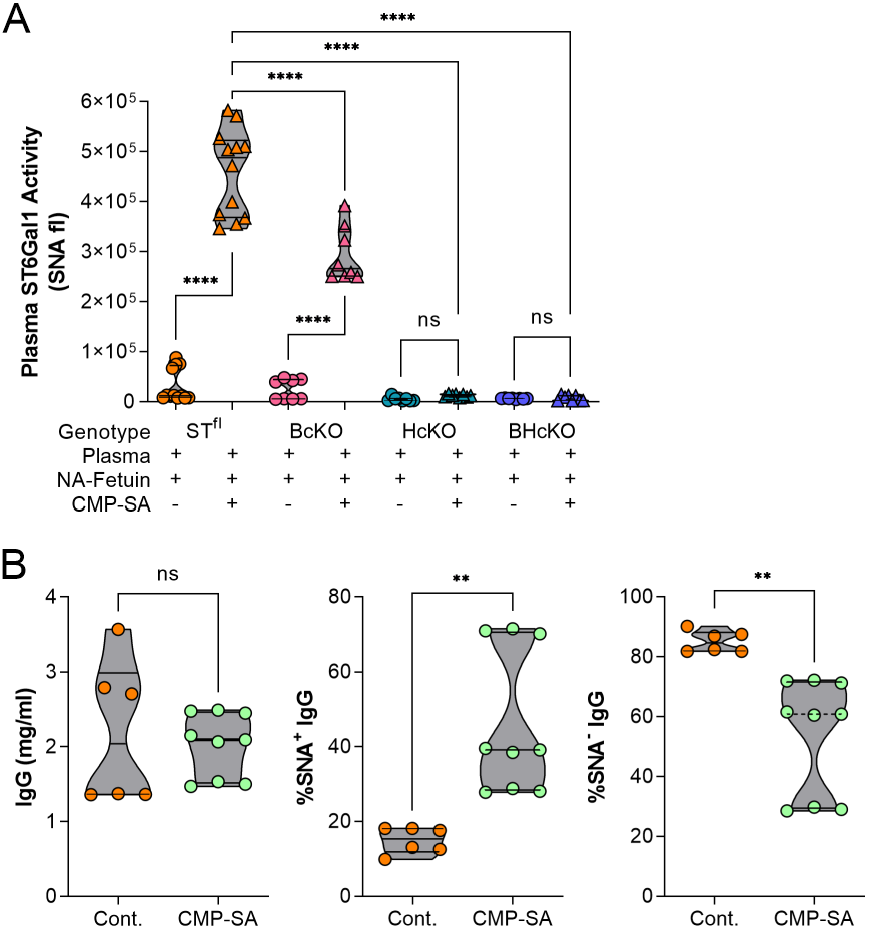
Plasma ST6Gal1 activity is lost in HcKO and BHcKO mice. (A) Addition of α2,6 sialic acids to plate-bound neuraminidase (NA)-treated fetuin with plasma from WT, BcKO, HcKO, and BHcKO mice, and with or without added CMP-sialic acid nucleotide-sugar donor, showing a loss of detectable activity in HcKO and BHcKO plasma. N=12 (B) I.v. injection of CMP-SA did not alter the total IgG concentration, but led to an increase in α2,6-sialylation in WT mice, confirming the function of plasma-localized ST6Gal1. N=6-9

Next, we analyzed the overall IgG titer (Fig 6A) and IgG subclass distribution (Fig 6B) in the WT, BcKO, HcKO, and BHcKO strains with and without immunization. In contrast to the data in Figure 2 in which IgG N-glycans were analyzed 2 weeks after the final immunization booster, we chose to perform the analysis at 8 weeks after the last injection to ensure all animals had returned to homeostasis following immune activation. In general, all three knockout strains had a slightly reduced IgG concentration following immunization. With the exception of the BcKO, IgG subclass distribution was also not substantially different from WT. The BcKO showed an expansion of IgG2a and IgG2b with a concomitant decrease in IgG1. More importantly, we found using mass spectrometry that the degree of sialylation and the relative distribution of sialic acid linkages was indistinguishable among all four strains (Fig 6C and 6D). A further breakdown of the dominant sialylated N-glycans revealed similar results to the WT and HcKO mice at two weeks in which singly sialylated species dominated (Fig 2), although there was an increase in dual sialylated species at this later time point (compare Fig 2E and 6E). These findings suggest that sialylation increases over time following a return to homeostasis (Fig 6E and 6F). These data reveal that both circulatory/hepatocyte as well as B cell ST6Gal1 are dispensable and do not compensate for each other.

**Figure 6.**
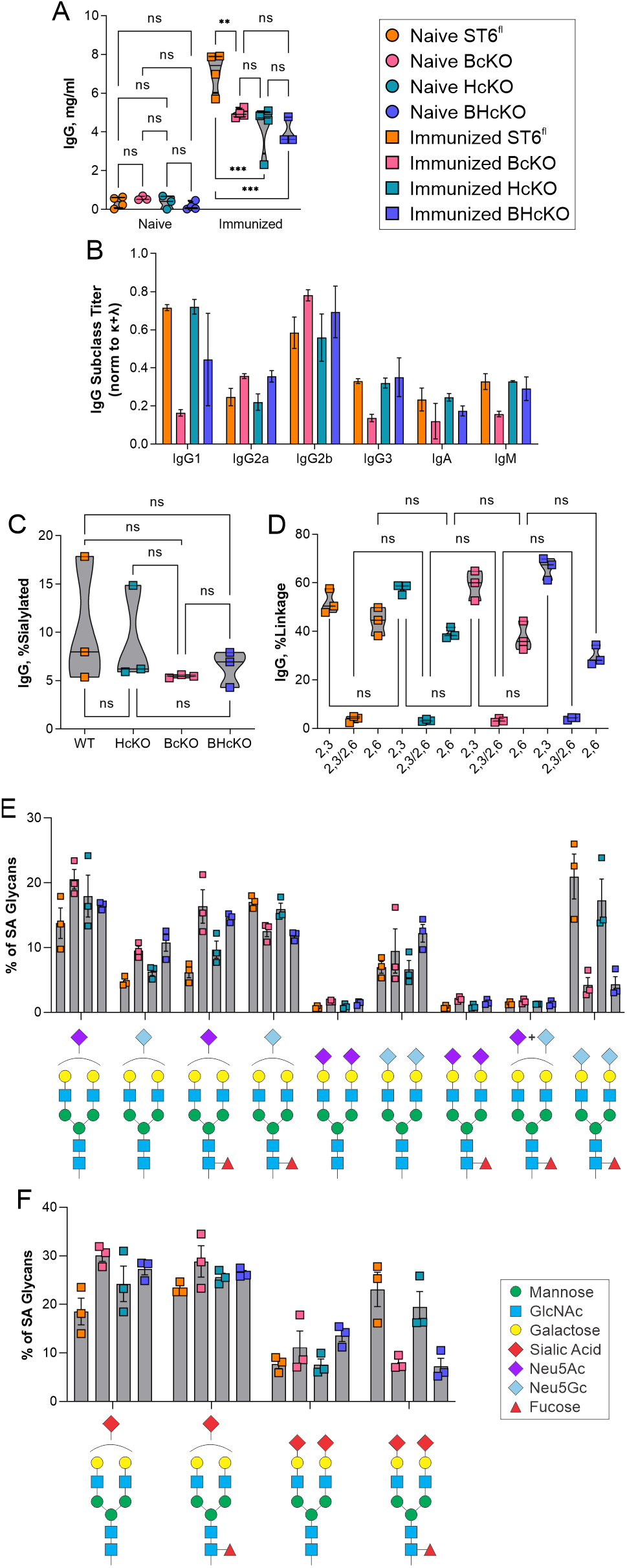
BHcKO IgG glycans mirror WT glycans. (A) IgG titers in immunized and non-immunized mice revealed slightly lower overall IgG titers in all three ST6Gal1 knockout mice. N=4 (B) With the exception of the BcKO strain, the Ig class and subclass distribution is essentially unchanged across strains. BcKO mice showed a decrease in IgG1 and IgG3 and an increase in IgG2a and IgG2b. N=3 (C) Based on mass spectrometry, overall IgG sialylation of IgG glycans 8 weeks post immunization was indistinguishable between strains. N=3 (D) Likewise, the distribution of α2,3 and α2,6 linkages were indistinguishable. (E and F) A breakdown of specific sialylated glycan species found showed that the general distribution of mono-sialylated, di-sialylated, Neu5Ac, Neu5Gc, and fucosylation is the same in all strains of mice, although G2FS2 structures, particularly with two Neu5Gc residues were higher in WT and HcKO compared to BcKO and BHcKO samples. N=3

Finally, the mass spectrometry data at 2 and 8 weeks post final immunization booster seemed to indicate a progressive sialylation process during the return to homeostasis after immunization/exposure (compare Fig 2F and 6F). Thus, we first analyzed each of the main sialylated species independently based on the distribution of α2,3 and α2,6 linkages. Among the mono-sialylated N-glycans, we found that there are shifts in the linkage distribution at different time points, but that this appears to be influenced by the strain (Fig 7A). For example, in the G2S1 N-glycan containing Neu5Ac, both WT and HcKO strains showed low initial α2,6 sialylation at 2 weeks, but that increased over time to yield equivalent α2,3 and α2,6 distributions by week 8. The equivalent distribution was not seen in BcKO or BHcKO mice at 8 weeks. Similarly, among the di-sialylated species, there was some change in the distributions, depending on the strain (Fig 7B). In general, we found that the relative amount of α2,6 linked sialic acids increased over time, and this correlated with an increase in di-sialylated N-glycans with increasing time after the last immunization in both WT and HcKO mice (Fig 8).

**Figure 7.**
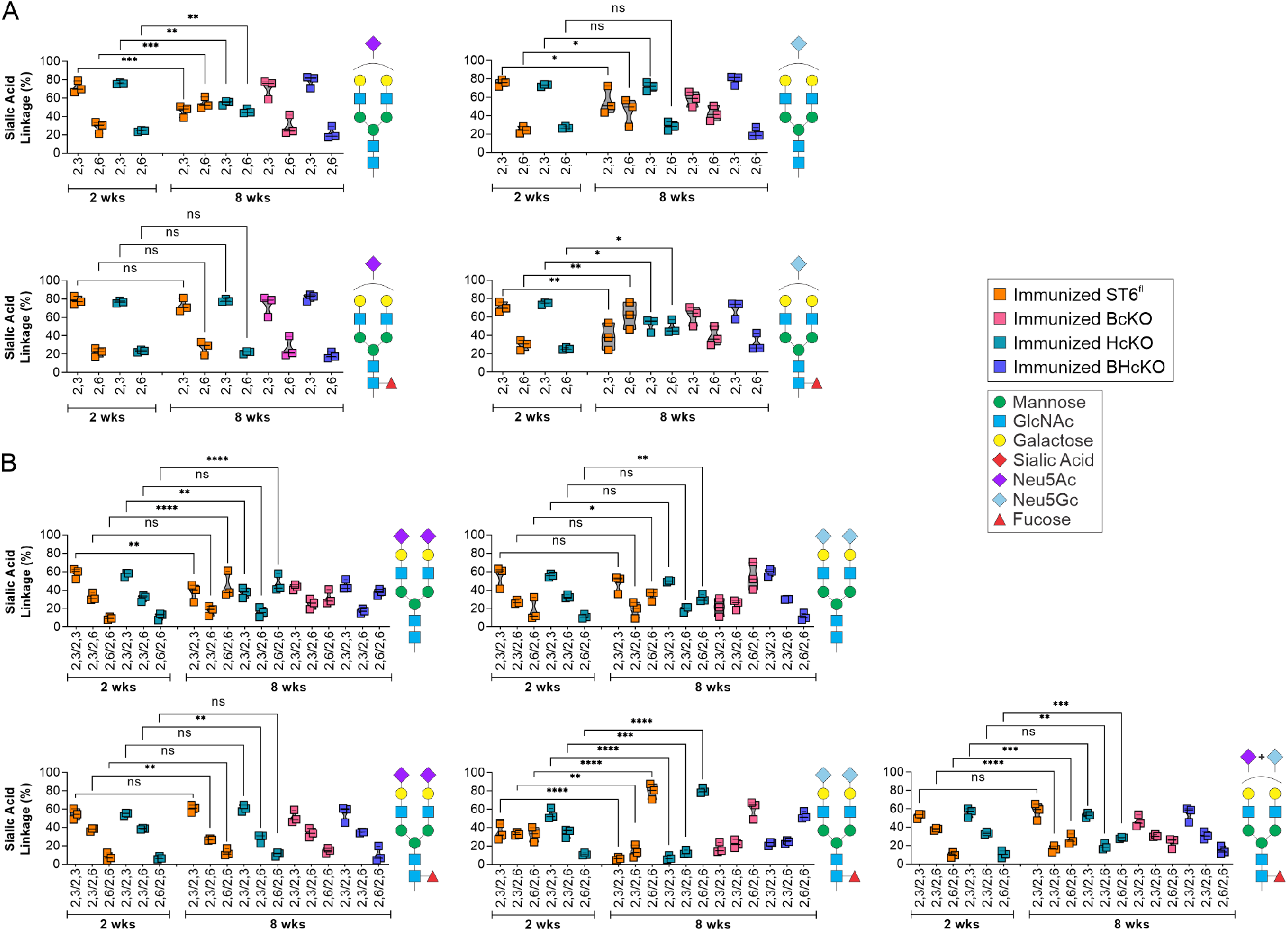
IgG sialylation linkage distribution is distinct for each sialylated species. (A) Comparison of the distribution of α2,3 and α2,6 linkages on a singly sialylated glycan species-by-species level at 2 weeks of WT and HcKO mice and 8 weeks of WT, HcKO, BcKO, and BHcKO mice. (B) Comparison of the distribution of α2,3 and α2,6 linkages on a dually sialylated glycan species-by-species level at 2 weeks of WT and HcKO mice and 8 weeks of WT, HcKO, BcKO, and BHcKO mice. The data revealed that linkage distributions change differentially among sialylated species. All data N=3

**Figure 8.**
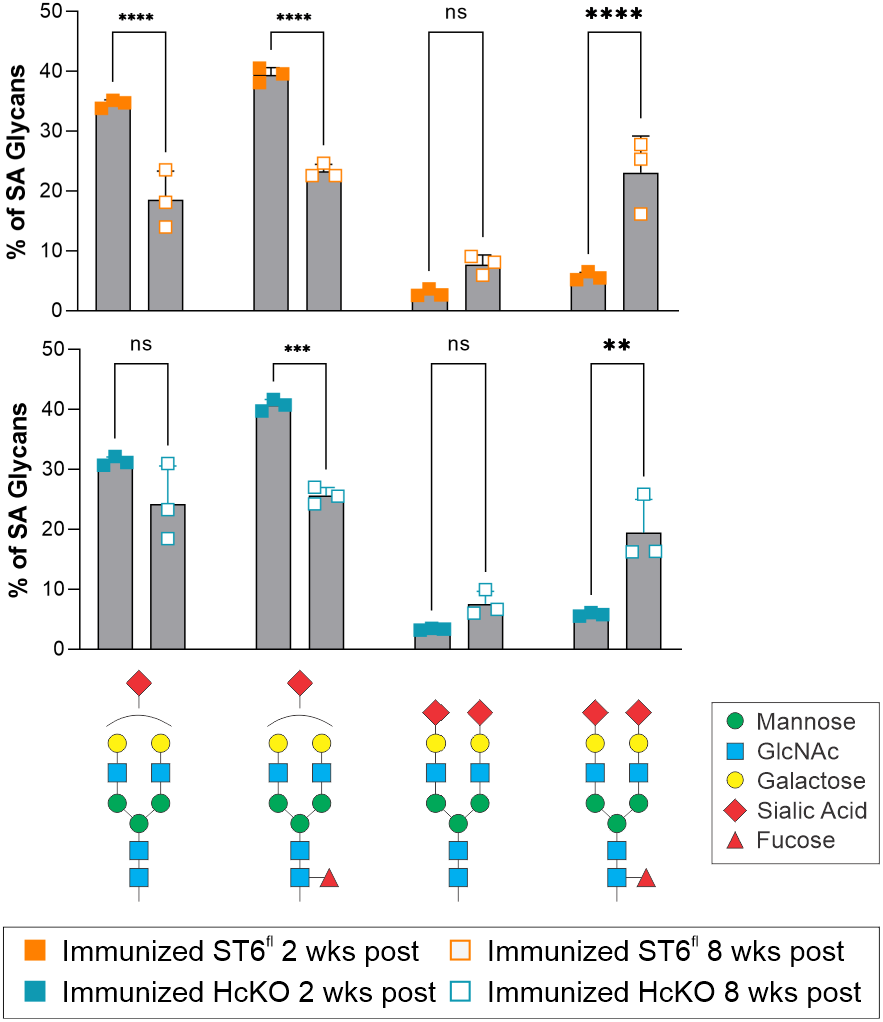
IgG sialylation increases over time following immunization. Time course comparison of the main singly and dually sialylated glycan species in WT and HcKO mice at 2 and 8 weeks post final immunization boost, which indicated a shift towards structures containing two sialic acids at a longer time point after the most recent exposure to antigen. N=3

## Discussion

Here we demonstrate, as might be expected, that α2,6-sialylation of IgG by ST6Gal1 arises by the action of ST6Gal1, as documented using germ-line deletion of ST6Gal1. Unexpectedly our prior studies had shown that such sialylation of IgG does not arise by the action of ST6Gal1 in B cells, although the loss of B cell ST6Gal1 led to a loss of cell surface α2,6-sialylation and replacement by terminal galactose (16). These prior studies led us to test the hypothesis that IgG α2,6-sialylation might arise through the action of liver-derived soluble ST6Gal1 in the plasma. However, our results demonstrate that neither the ST6Gal1 in plasma or in B cells is chiefly responsible for IgG α2,6-sialylation.

The plasma-derived ST6Gal1 is known to be released from the liver through the action of the protease BACE-1 (21,22). The notion that IgG sialylation might involve plasma ST6Gal1 originated in studies utilizing a mouse with the P1 region of the ST6Gal1 promoter ablated (23). The ΔP1 mouse has a documented reduction, albeit not an ablation, of plasma ST6Gal1 activity along with a reduction in IgG sialylation (19,23). We considered that the creation of the BcKO mouse and the observation that IgG sialylation was unchanged (19) seemed to support this model. Through the creation of the combined HcKO and BHcKO mice, our results here demonstrate that plasma ST6Gal1 is not necessary for IgG sialylation. Complete loss of plasma ST6Gal1 activity, ablation of BACE-1, and a combination knockout in hepatocytes and B cells collectively demonstrate that IgG sialylation occurs elsewhere, but certainly not in either B cells nor plasma.

The lack of plasma ST6Gal1 making an impact on glycans within the circulation is further supported by the fate of the B cells in the BcKO mouse (19). These cells have a dramatically reduced surface α2,6-sialylation, yet are in direct contact with ST6Gal1 in plasma just like IgG. We were puzzled by the apparent contradiction, that whereas B cell surfaces lacked α2,6-sialylation, their secreted IgG was 2,6-sialylated. The current HcKO mouse supports the interpretation that plasma ST6Gal1, while functional, cannot functionally sialylate B cells lacking endogenous ST6Gal1. This is most likely due to the lack of CMP-sialic acid required as a donor for the sialylation reaction under most conditions.

Another key uncertainty has been why the ΔP1 mouse does not phenocopy the HcKO strain. The original description of the ΔP1 mouse exhibited a modest reduction in circulatory ST6Gal1 protein and activity using *in vitro* assays, likely due to the fact that expression of the enzyme is incompletely knocked down. Moreover, since the creation of the ΔP1 mouse, it has become clear that these animals carry interesting hematopoietic phenotypes (29–31) that are also not phenocopied in the HcKO mice (25,26). We reason that the manipulation of the P1 region of the promoter has an impact beyond the liver and circulatory activity of ST6Gal1, and that it is likely involved with expression of ST6Gal1 in multiple hematopoietic cell lineages. Given that the HcKO mouse lacks detectable ST6Gal1 activity in plasma, we believe that non-liver reductions in expression are the source of the phenotypes described using the ΔP1 mouse, and not changes in plasma ST6Gal1.

Beyond the continuing puzzle regarding the non-plasma and non-B cell location of IgG sialylation, another observation herein is the apparent high proportion of α2,3-linkages among sialylated N-glycans found on IgG in these mice. Work citing α2,3 sialylation of IgG is extremely limited, although α2,3- and α2,6-sialylation of murine IgG was recently documented, as well as strain differences. Unfortunately, a great number of studies on IgG glycosylation does not address the specific nature of sialic acid linkages. As such, much of the discussion and analysis of IgG sialylation may be, to some degree, problematic. The effects of α2,3-vs α2,6-sialylation on the activities of IgG are not well documented. Thus, it is likewise not clear whether α2,3 sialylation of IgG affects its affinity to the Fcγ receptor in the same way as α2,6-sialylation, nor is it known how α2,3-sialylation might impact IgG structure. In this regard, while IgG glycosylation invariably occurs on the Fc portion at N297, it is known that additional N-glycans might be present on a subset of IgG through N-glycosylation of the Fab domain. However, this would not affect our overall interpretation, as we analyzed total N-glycans and their total sialylation, in which α2,6-sialylation could of course impact both types of N-glycans.

Finally, we found that the sialylation of IgG N-glycans changes as a function of distance in time from the most recent exposure or immunization. There have been proposals as to the possibility of programming the glycoform of specific clones of antibodies *in vivo* via vaccination (32), but our data suggest that IgG glycosylation is perhaps not stable enough for this to work as a therapeutic approach, at least in the murine system, since the degree of sialylation as well as the proportion of singly or doubly sialylated species changes longitudinally after immunization.

In summary, our data now allows us to reject the model that IgG sialylation occurs within the circulatory microenvironment through plasma-localized ST6Gal1, as well as documenting a significant degree of α2,3 sialylation on IgG N-glycans in mice, and an unstable degree of sialylation over time following immunization. The findings point to an unidentified anatomical location or cell type in which IgG is sialylated. A leading candidate may be platelets, given their expression of ST6Gal1 and demonstrated ability to release CMP-sialic acid (16,24), yet much more research is needed to clarify these mechanisms *in vivo*.

## Experimental procedures

### Animal care and use

All animal work was approved by the institutional animal care and use committee (IACUC) of Case Western Reserve University. Mice were procured from Jackson Laboratories and kept/bred in the CWRU ARC in accordance with IACUC guidelines. ST6Gal1^f/f^ mice (stock 006901) and Alb-Cre mice (stock 003574) were crossed to produce the HcKO line, as we document in our previous manuscript (26). Wherever the phrases “wild type” or “parental” are used, they refer to mice of the ST6Gal1^f/f^ background, which carry no phenotype, with the exception of the control mice for the BACE-1 study which were C57Bl/6J mice directly housed by Jackson Laboratories (JAX). In that case, the WT and BACE-1 knockout mouse serum was harvested by and purchased directly from JAX. BcKO mice were produced by crossing the ST6Gal1^f/f^ mice with CD19-Cre mice (stock 006785), and were used as experimental when homozygous on the allele and heterozygous on the CD19-Cre allele, as described (16). BHcKO mice were combined by crossing homozygous breeder of the BcKO and HcKO line, resulting in ST6Gal1^f/f^, Alb-Cre^+/-^, CD19-Cre^+/-^ mice.

### Animal harvest

For harvest, mice were euthanized using CO_2_ following standard procedures. Blood was drawn via cardiac puncture. Plasma was separated from blood by spinning at 2000 x g for 15 minutes, and aspirating the liquid phase.

### Flow cytometry

Flow cytometry was performed on cells isolated freshly *ex vivo* from mice, as described elsewhere (16). Liver lobes were diced and digested in 2.5 mg/ml collagenase (cat) in 3 % FBS for 45 minutes at 37 °C. Tissues were diced and mashed through 70 um nylon filters in 10 ml of PBS. Cells were spun down, then RBC lysis was performed using pharmlyze (BD Biosciences, 555899) at 4 °C for 5 minutes, then quenched with PBS. Cells were blocked for 30 minutes in Carbohydrate Free Blocking Solution (Vector) and stained with the following reagents: SNA-FITC (Vector), CD19-APC (Biolegend). Flow cytometry was run on an Accuri C7. Data was analyzed using FlowJo.

### Histology

Histology embedding and sectioning was performed by the CWRU Cancer Center Histology Services Core as previously described (16). Tissues for histological analysis were resected from mice and fixed in 10% formalin solution. Tissue blocks were embedded in paraffin wax, sectioned, and stained with H&E. For confocal analysis, freshly cut sections were de-parafinized in xylenes and rehydrated. The tissue was blocked using carbohydrate free blocking solution (Vector) and stained with SNA-FITC (Vector) at 0.5 μg/ml. Autofluorescence was diminished using TrueVIEW autofluorescence Quenching Kit (Vector) and the slides were mounted using VECTASHILED HardSet Antifade Mounting Medium (Vector). Confocal imaging was performed on a SP5 Laser Scanning Confocal Microscope (Leica).

### IgG purification

IgG was purified by separation over a HiTrap Protein A HP Antibody Purification Column (GE Life Sciences, 17040203) fitted to a GE Life Sciences Akta Purifier 10 HPLC, according to manufacturer instructions. IgG was tittered using an IgG ELISA kit (Bethyl, E90-131) according to manufacturer instructions.

### Mass spectrometry

Mass spectrometry N-glycan profile and sialic acid linkage analysis was performed on purified IgG essentially as described previously following DMT-MM and permethylation derivatization (28). IgG samples were lyophilized and first incubated with 2 mg/mL 1,4-dithiothreitol (Millipore-Sigma) 600 mM Tris buffer at pH 8.5 for 1 hour at 50 °C, then 12 mg/mL iodoacetamide (Millipore-Sigma) also in 600 mM Tris at pH 8.5 for 1 hour in the dark. Following dialysis against 50 mM ammonium bicarbonate, samples were dried by lyophilization and resuspended in 50 mM ammonium bicarbonate buffer containing 25 μg of TPCK-treated trypsin (Millipore-Sigma) and incubated overnight at 37 °C. The trypsin-digested samples were cleaned using a 50 mg C18 Sep-Pak column (Waters). The lyophilized samples were then resuspended in 50 mM ammonium bicarbonate buffer and digested with PNGaseF (New England Biolabs) for a total of 20 hours. The PNGaseF-released N-glycans were isolated on a C18 Sep-Pak column, evenly split into two fractions and dried.

One of the two dried N-glycan aliquots for each sample was dissolved in a 1 mL slurry solution of NaOH in DMSO. 500 μL of iodomethane (Millipore-Sigma) were added and the samples were mixed vigorously at room temperature for 30 minutes. 1 mL of MilliQ water was then added to stop the reaction. 1 mL of chloroform and 3 mL of MilliQ water were added, vortexed thoroughly, and centrifuged (5000 rpm, 30 sec). The aqueous top layer was discarded and 3 mL of MilliQ water was added to wash the chloroform phase by vortexing and centrifuging as before. This was repeated one additional time before drying the chloroform fraction with a speedvac. The dried permethylated N-glycans were cleaned on a C18 Sep-Pak column and eluted with 3 mL of 50 % acetonitrile. The eluted fraction was lyophilized prior to MS analysis.

For the DMT-MM (4-(4,6-dimethoxy-1,3,5-triazin-2yl)-4-methylmorpholinium chloride) treatment, a solution of 0.5 M of DMT-MM (Millipore-Sigma) was prepared in 500 mM of NH_4_Cl, pH 6.5. 10 μL of the DMT-MM solution were added to the second aliquot of each N-glycan sample and incubated at 60 °C for 15 hours. Samples were cleaned on a C18 Sep-Pak column, lyophilized, and permethylated as before.

Permethylated N-glycans +/- DMT-MM derivatization were dissolved in 10 μL of 75 % methanol, from which 1 μL was mixed with 1 μL 2,5-dihydroxybenzoic acid (DHB) (5 mg/mL in 50% acetonitrile with 0.1% trifluoroacetic acid) and spotted on a MALDI polished steel target plate (Bruker Daltonics). MS data were acquired on a Bruker UltraFlex II MALDI-TOF Mass Spectrometer instrument. The reflective positive mode was used, and data were recorded between 500 and 6000 m/z. For each MS N-glycan profile, the aggregation of 20,000 laser shots or more were considered for data extraction. Only MS signals matching an N-glycan composition were considered for further analysis. Subsequent MS post-data acquisition analysis was made using mMass (33).

### Lectin ELISA

High throughput, multiplexed lectin ELISA was performed on whole plasma samples, as has been published (27,34). Briefly, plasma was diluted to 0.5 μg/ml in carbonate coating buffer (100 mM NaHCO_3_, 30 mM NaCO_3_, pH 9.5), pipetted into a 384 well black ELISA plate, and incubated overnight at 4 °C. The plate was blocked with carbohydrate free blocking solution (Vector, SP-5040) for 1 hour at room temperature. Biotinylated lectins were diluted as previously described and incubated on the plate for 1 hour at room temperature. Signal was detected using streptavidin-Eu (Perkin Elmer 1244-360) and enhancement solution (Perkin Elmer, 4001-0010) measured in a Victor V3 1420 multilabel plate reader.

### Lectin chromatography

Lectin chromatography of IgG was performed as previously described (16). Purified IgG was fractionated using SNA-agarose (Vector) into SNA^-^ fraction, then washed 3 times (10 mM HEPES, 0.1 % TX-100, pH 7.5) and an SNA^+^ fraction was eluted (100 mM glycine, 100 mM sodium acetate, 5 mM MgCl_2_, pH 4.5). IgG concentration was determined as detailed above.

### ST6Gal1 activity

A 96 well ELISA plate was coated with bovine fetuin (5 μg/mL; Sigma) in carbonate buffer overnight, then blocked for 1 hour at room temperature with Carbohydrate-Free blocking solution (Vector Labs), and treated with 0.005 U of neuraminidase from *C. perfringens* (Sigma) per well in 1 mM Na_2_HPO_4_, 1 mM K_2_HPO_4_, and 10 mM KCl at pH 6.0 for 1 hour at 37 °C. Using harvested mouse plasma, an enzymatic reaction was performed at 37 °C for 3 hours with or without 100 μM CMP-sialic acid donor nucleotide-sugar. Added α2,6-linked sialic acids were probed using SNA-biotin (0.4 μg/mL), Eu-conjugated streptavidin (Perkin-Elmer) at 0.1 μg/mL, and detected on a Victor V3 multilabel plate reader as previously published (16).

### CMP-SA injection

Mice were injecting in the tail vein with 200 μg of CMP-SA suspended in PBS and plasma harvested after 24 hours.

### Data Analysis

Data was analyzed using Microsoft Excel or R and plotted in GraphPad Prism. The R function “prcomp” was used to calculate principal component analysis. PCA was then blotted using the “fviz_pca_biplot” function. Statistical testing was done by student’s T test: *, p < 0.01: **. p < 0.001: ***.

## Acknowledgements

The authors wish to acknowledge Dr. Mark B. Jones for scientific input in the early phases of this project, and for the BACE-1 knockout IgG analysis. Funding for this work was through grants to BAC from the NIH (GM115234 and AI154899) and to JYZ and DMO from the NIH (AI089474).

## Conflict of interest

The authors declare that they have no financial conflicts of interest with the contents of this article.

## Author Contributions

DMO, experimental design, data collection and analysis, manuscript writing; JYZ and LMG, data acquisition; SDL and RDC, mass spectrometry design, data collection and analysis, manuscript editing; BAC, experimental design, data analysis, manuscript writing, funding.

## Abbreviations

(DMT-MM): 4-(4,6-dimethoxy-1,3,5-triazin-2yl)-4-methylmorpholinium chloride
(BcKO): B cell-specific conditional knockout of ST6Gal1
(CMP-sialic acid): cytidine monophosphate N-acetylneuraminic acid
(HcKO): hepatocyte-specific conditional knockout of ST6Gal1
(IVIg): intravenous immunoglobulin therapy
(BHcKO): mouse lacking ST6Gal1 in both the hepatocyte and B cell compartments
(PCA): principle component analysis
(RA): rheumatoid arthritis
(WT): wild type

## Notes

### Competing Interest Statement

The authors have declared no competing interest.

